# Conserved tryptophan mutation disrupts structure and function of immunoglobulin domain revealing unusual tyrosine fluorescence

**DOI:** 10.1101/2020.06.22.165159

**Authors:** Ravi Vattepu, Rachel A. Klausmeyer, Allan Ayella, Rahul Yadav, Joseph T. Dille, Stan V. Saiz, Moriah R. Beck

## Abstract

Immunoglobulin (Ig) domains are the most prevalent protein domain structure and share a highly conserved folding pattern; however, this structural family of proteins is also the most diverse in terms of biological roles and tissue expression. Ig domains vary significantly in amino acid sequence but share a highly conserved tryptophan in the hydrophobic core of this beta-stranded protein. Palladin is an actin binding and bundling protein that has five Ig domains and plays an important role in normal cell adhesion and motility. Mutation of the core tryptophan in one Ig domain of palladin has been identified in a pancreatic cancer cell line, suggesting a crucial role for this sole tryptophan in palladin Ig domain structure, stability, and function. We found that actin binding and bundling was not completely abolished with removal of this tryptophan despite a partially unfolded structure and significantly reduced stability of the mutant Ig domain as shown by circular dichroism investigations. In addition, this mutant palladin domain displays a tryptophan-like fluorescence attributed to an anomalous tyrosine emission at 345 nm. Our results indicate that this emission originates from a tyrosinate that may be formed in the excited ground state by proton transfer to a nearby glutamyl residue. Furthermore, this study emphasizes the importance of tryptophan in protein structural stability and illustrates how tyrosinate emission contributions may be overlooked during the interpretation of the fluorescence properties of proteins.

**SHORT ABSTRACT:** This study explores the functional and structural consequences of a point mutation in palladin, an Ig domain protein first identified in a pancreatic tumor cancer cell line. While exploring the consequences of mutating this conserved tryptophan in the hydrophobic core of the most prevalent domain structure found in proteins, an anomalous tyrosine fluorescence phenomenon was exposed.

## 1 INTRODUCTION

The Ig domain fold is found in a diverse set of proteins that vary significantly in amino acid sequence, tissue distribution, and biological roles. This diverse set of proteins adopt a similar fold for stability and to mediate protein-protein interactions. Typical Ig-like domains consist of 100-120 amino acids that fold into the characteristic β-sandwich fold; however, the number of Ig domains in each protein varies considerably.^1^ Generally, immunoglobulin domains are composed of two beta sheets that are both right-hand twisted and stack on top of one another enclosing the hydrophobic core that forms around conserved aromatic amino acids tryptophan and tyrosine (Figure 1a).^1^ Ig domains are known to mediate a variety of protein-protein interactions and play significant roles in cell-cell adhesion and signaling both inside the cell and at the cell-surface.

**Figure 1.**
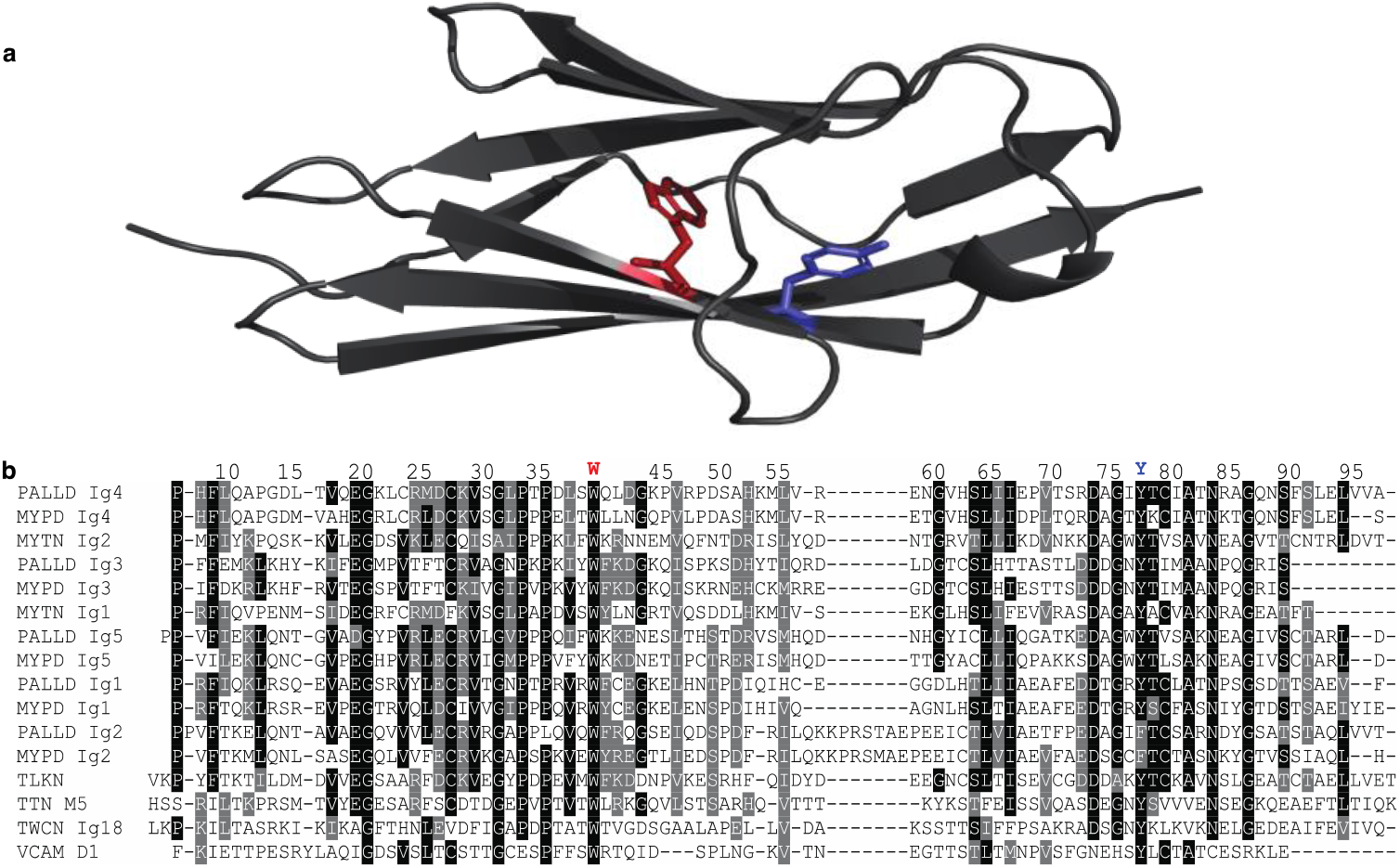
Structure and sequence alignment of palladin Ig4 domain. (a) Ribbon representation of the Ig4 domain (PDB ID: 2DM3), where conserved tryptophan and tyrosine residues are highlighted as sticks: W40 (red) and Y78 (blue). (b) Sequence alignment of Ig4 with other Ig domain containing proteins.

Palladin, myotilin, and myopalladin form a small family of intracellular actin regulatory proteins that each contain between two to five tandem Ig domains.^2; 3^ Palladin mediates several different protein-protein interactions through one of two polyproline-rich regions as well as the N-terminal and the C-terminal Ig domains. The Ig3 domain of palladin is directly involved in actin binding, polymerization, and cross-linking; however, the Ig4 domain increases these functions synergistically in the tandem Ig3-4 domain.^4–6^ We have shown that actin-induced dimerization of both the Ig3 and Ig3-4 domains of palladin promotes actin bundling; and, furthermore, the extent of dimerization reflects the degree of actin crosslinking.^6^ The Ig3 and Ig4 domains of palladin and myopalladin are homologous to the Ig1 and Ig2 domains of myotilin. Interestingly, the tryptophan located in the C strand of the Ig domain is absolutely conserved across all of the Ig domains in the palladin family and many other Ig domain containing proteins as shown in Figure 1b.

Increased expression of palladin in both pancreatic and breast cancer cell lines suggests a role for palladin in the invasive motility associated with metastasis.^7^ Additional evidence linking palladin to metastatic cancer comes from a genetic comparison of different sporadic pancreatic cancer cells lines that revealed that the highly metastatic pancreatic tumor (PaTu2) cell line has one specific mutation that changes the highly conserved tryptophan to cysteine (W40C for Ig4 domain numbering or W1168C in the full-length, 90 kDa palladin) in the Ig4 domain of palladin.^8^ When COS-7 cells were transfected with a PaTu2 mutant version of palladin, the expression pattern was different than in cells expressing WT palladin.^8^ Namely, low levels of PaTu2 palladin expression were associated with weak actin filaments and actin bundle formation as compared to WT. In addition, WT palladin is involved in the contraction and aggregation of stress fibers at higher expression, but PaTu2 forms smaller aggregates that lack filamentous actin.^8^

Since the mutated tryptophan in PaTu2 Ig4 is a consistent feature of Ig domains in palladin, as well as Ig domains in general, we hypothesize that it is necessary for proper domain folding and therefore function. Although this mutation has not been observed in clinical samples of primary tumors, understanding how this mutation alters the structure and function of the palladin Ig4 domain will increase our understanding of the broader role of palladin in promoting metastasis and allow us to examine the role of this conserved tryptophan in immunoglobulin domain folding and stability. To understand the influence of this mutant on palladin domain function and structure, we have employed a series of *in vitro* experiments exploring the protein function, dynamics and folding behavior.

Our studies in the mutant palladin Ig domain lacking the conserved tryptophan led to the surprising observation of tyrosine fluorescence resembling a typical tryptophan. We compare our experimental spectroscopic results with previously observed anomalous tyrosine fluorescence^9–14^ and present support for a mechanism similar to that proposed by Pundak and Roche^13^ that explains the fluorescent properties of tyrosine in the mutant palladin domain.

## 2 RESULTS

### 2.1 Effect of palladin PaTu2 mutation on actin binding versus bundling

Previous work has already established that the Ig4 domain of palladin does not bind or bundle actin, but does enhance actin bundling synergistically in the tandem Ig3-4 domain.^5^ To compare the wildtype Ig3-4 and Ig4 domains of palladin with the PaTu2 mutation, we genetically engineered a point mutation to replace the sole, conserved tryptophan in the Ig4 domain with a cysteine. We then determined how this mutation affects actin binding and bundling. Our actin co-sedimentation results provide clear evidence that there is only a slight change in the actin-binding affinity (WT Ig3-4 K_d_ = 2.9 μM and PaTu2 Ig3-4 K_d_= 6.0 μM) (Figures 2a and b), while there is a significant decrease in actin bundling for the mutant (Figures 2c and d). At increased concentrations of PaTu2 Ig3-4, the actin bundle formation is similar to wildtype Ig3-4; however, at lower concentrations, we observed a significant decrease for the PaTu2 mutant. Specifically, at the 0.5:1 and 1:1 ratios of actin:palladin Ig domain, the PaTu2 Ig3-4 mutant results in only ∼20 and ∼40 percent of actin in bundles, while WT Ig3-4 yields ∼55 and ∼95 percent actin bundle formation as shown in Figures 2c and d. These results reinforce the notion that the Ig4 domain is critical for actin bundling as mutations in the Ig4 domain primarily affect the actin bundling and not binding as shown in Figure 2.

**Figure 2.**
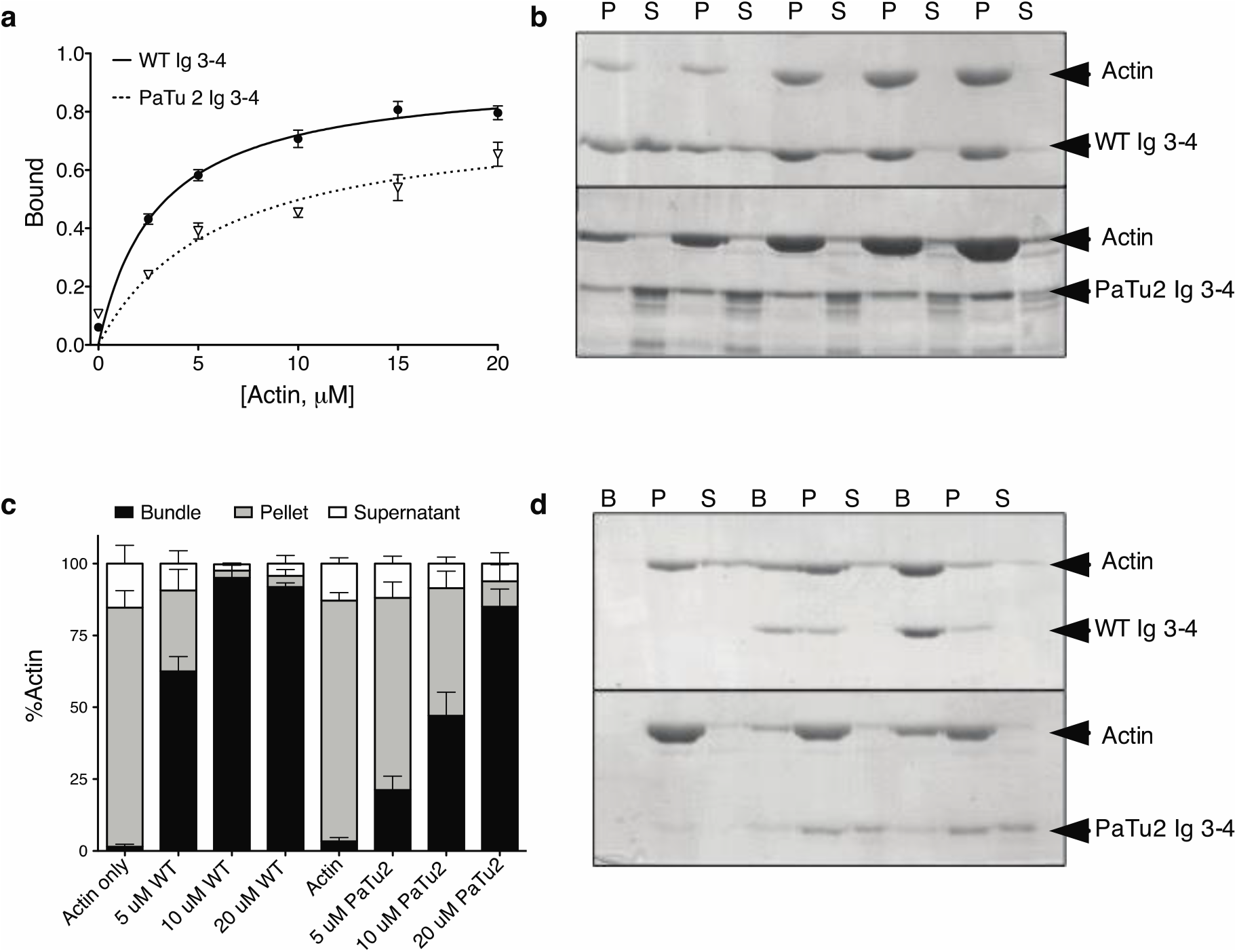
Comparison of actin-binding and bundling for WT and PaTu2 Ig3-4 domains of palladin. (a) Quantification of actin binding analyzed by densitometry of the 5 μM WT or PaTu2 Ig3-4 in pellet fractions plotted as a function of actin concentration. (b) Representative SDS-PAGE gel from one of three independent co-sedimentation assays. (c) Quantification of actin bundling analyzed by densitometry of the amount of actin present in the bundle, pellet, and supernatant fractions in the presence of WT or PaTu2 Ig3-4. (d) Representative SDS-PAGE gel from one of three independent bundling assays.

### 2.2 PaTu2 mutation alters secondary structure

We also employed circular dichroism (CD) spectroscopy to characterize the secondary structure, where the high β-strand content of immunoglobulin domains typically display a minimum at 217 nm and a zero intensity peak at 206 nm. We observed a peak minimum for each palladin Ig domain in the far UV CD spectra that differs significantly as shown in Figure 3. The peak minima for both PaTu2 mutant proteins (PaTu2 Ig4 and Ig3-4) are very broad and shifts to lower wavelengths with respect to WT. Furthermore, a peak maximum was observed at 206 nm only for WT Ig3-4, while WT Ig4 has a negative peak with a sharp transition. The PaTu2 mutants also display negative peaks, but also a much broader transition. Overall these spectral variations can be attributed to significant structural differences between WT and PaTu2 Ig4 domains.

**Figure 3.**
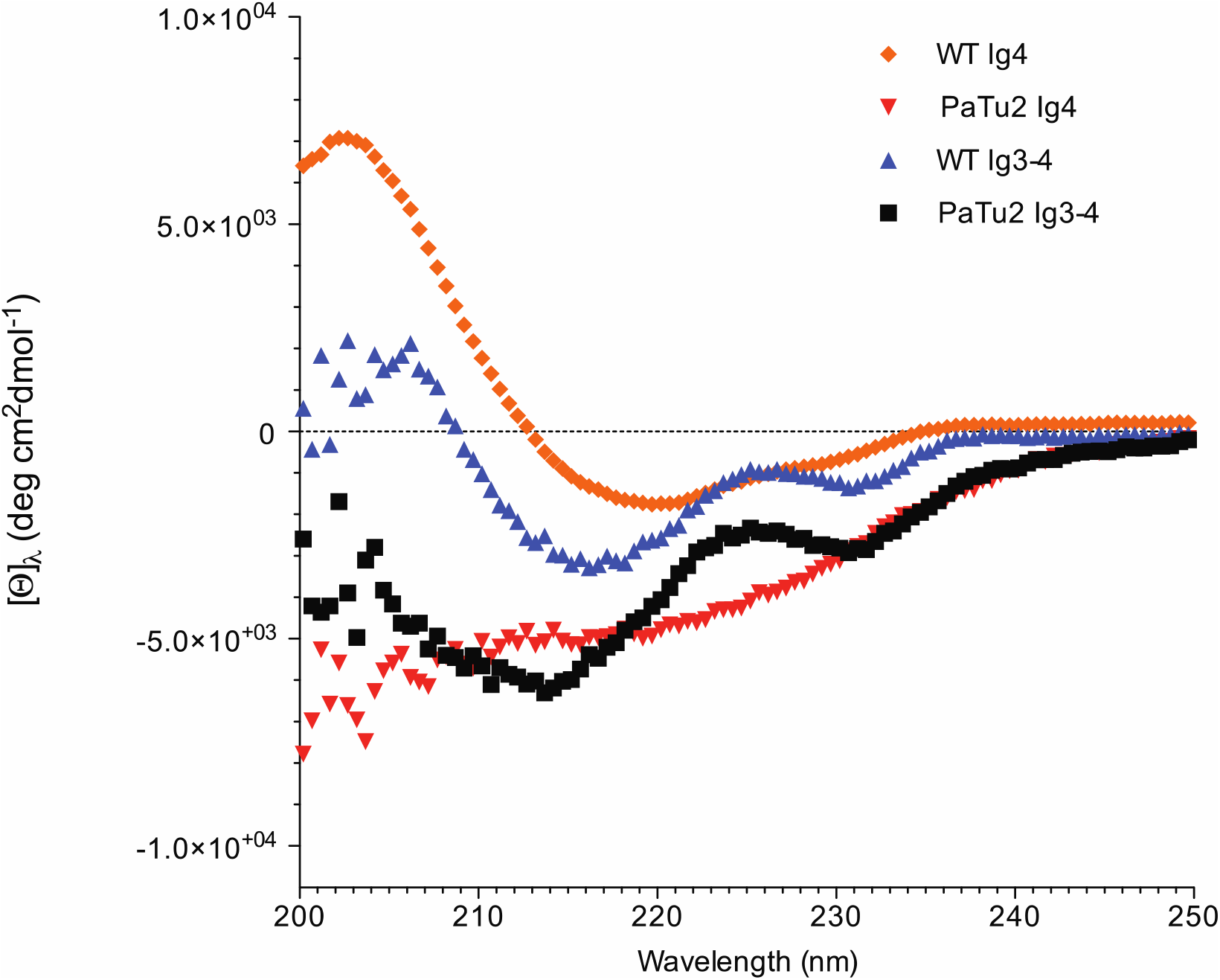
Palladin Ig domains secondary structure comparison using circular dichroism. Room temperature CD spectra of various 20 μM palladin domains.

### 2.3 PaTu2 mutation disrupts Ig4 domain folding

To determine the role of this conserved tryptophan in the structure of the Ig4 domain, we also collected ^1^H-^15^N heteronuclear single quantum correlation spectroscopy (HSQC) NMR spectra of PaTu2 Ig4 to compare with the WT Ig4 HSQC spectrum (Figure 4). We observed a significant decrease in the chemical shift dispersion in the ^1^H dimension of the PaTu2 Ig4 spectrum, implying that this mutation caused significant unfolding of the protein. This is further confirmed by overlaying the HSQC spectra of WT and PaTu2 Ig4, where significant differences in the chemical shift dispersion can easily be identified. These solution-state NMR results reveal that this mutation significantly disrupts the Ig domain folding and this is consistent with the fact that all Ig domains have a conserved, buried tryptophan that is thought to play a critical role in proper Ig domain folding.

**Figure 4.**
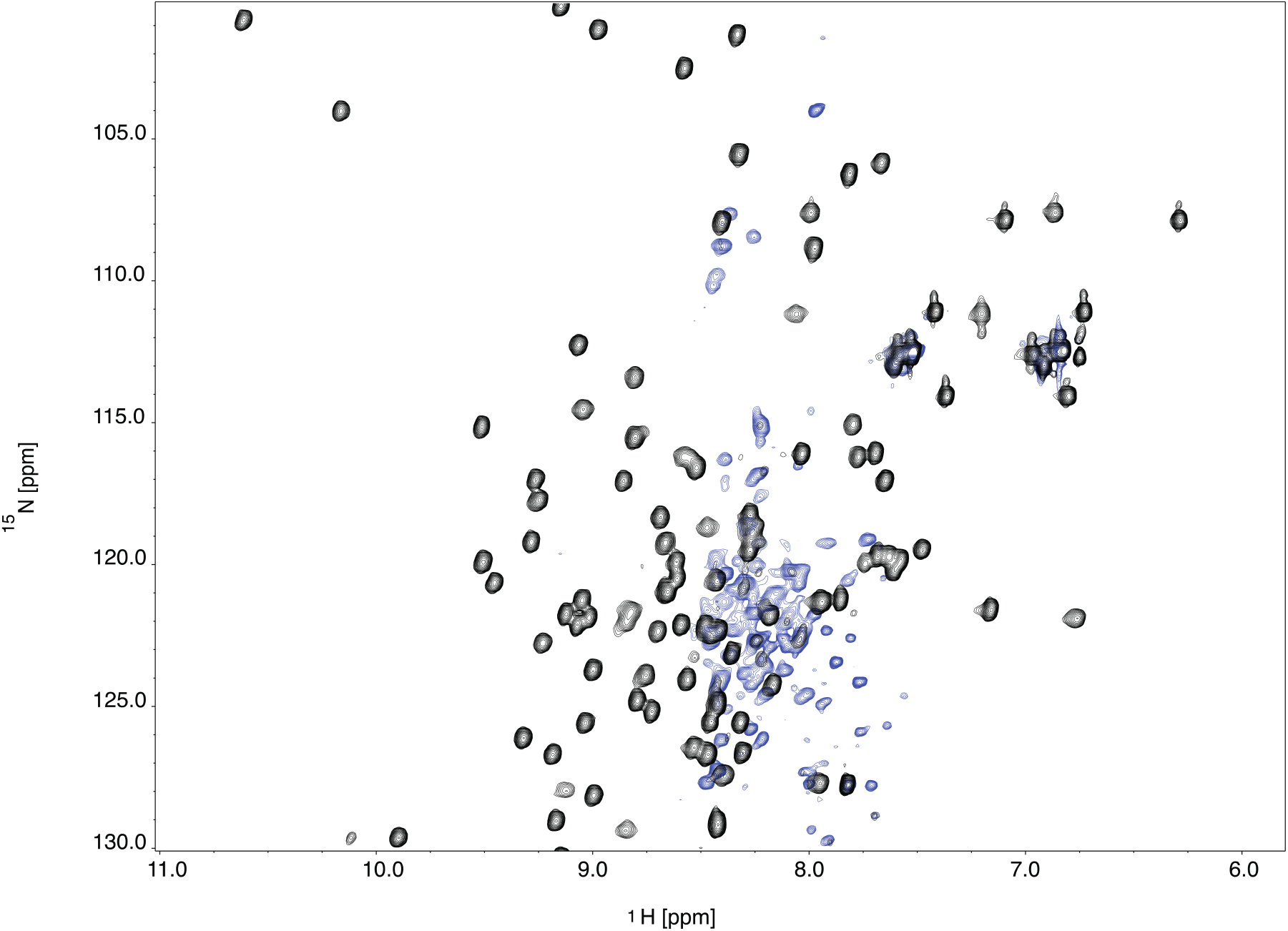
Protein NMR studies revealed that PaTu2 mutation disrupts Ig4 domain folding. Overlay of ^1^H–^15^N HSQC spectra of 0.2 mM WT Ig4 (black) and PaTu2 Ig4 (blue).

In line with this observation, we have also detected a significant difference in the bacterial expression pattern of the PaTu2 protein (results not shown). WT Ig4 is expressed successfully without the need for any solubility tag, while PaTu2 Ig4 is segregated to inclusion bodies. Purification of soluble, stable PaTu2 Ig4 required the addition of a maltose binding protein (MBP) tag. This lack of solubility for the PaTu2 mutant of Ig4 strengthens the hypothesis that the conserved tryptophan in the hydrophobic core influences the Ig domain folding.

### 2.4 Palladin Ig domain stability

To determine the effects of the PaTu2 mutation on protein stability, the fraction folded was calculated from each fluorescence emission spectra as a function of urea concentration as shown in Figure 5. The transition mid-point of each emission spectra is calculated by fitting to equation 1, yielding the transition mid-points (C_m_) listed in Table 1. The stability of the PaTu2 Ig4 mutant is significantly decreased as compared to WT which suggests that the W40C mutation affects both structure and stability. Interestingly, the PaTu2 Ig3-4 exhibits two transitions during unfolding, while WT Ig3-4 displays only a single transition. Three transition states (native, intermediate and unfolded) were calculated using equation 2 for PaTu2 Ig3-4 (see Methods section). We surmise that the first transition results from PaTu2 Ig4 domain unfolding with lower transition midpoint, while the second transition reflects the unfolding of the more stable Ig3 domain with a higher transition midpoint. We have previously shown, using chemical crosslinking experiments, that the Ig3 and Ig4 domains interact with each other and upon actin binding these two domains undergo a conformational change that appears to separate the domains from each other.^6^ In these unfolding experiments, the observed single transition of WT Ig3-4 may also be influenced by the fact that these two domains interact with each other and therefore unfold as one. Conversely, the PaTu2 mutation disrupts Ig4 domain folding which could potentially interfere with the interactions between the two domains. This interference could cause the two domains to unfold independently giving rise to two unfolding transitions.

**Table 1.**
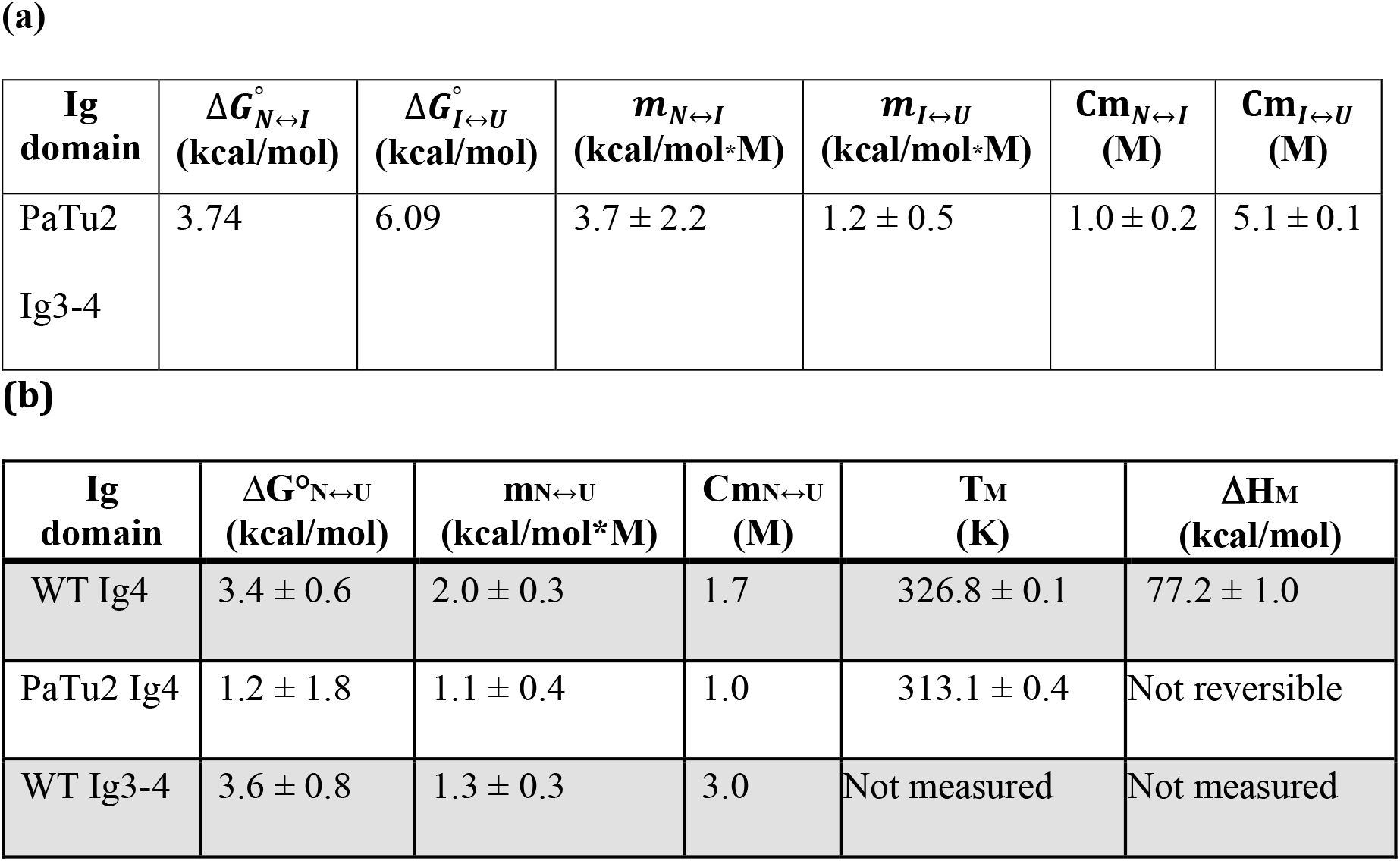
Thermodynamic parameters for denaturation curves of Ig4 and Ig3-4. **(a)** Thermodynamic parameters of the Ig3-4 PaTu2 from urea denaturation curve determined using the three-state model. **(b**) Thermodynamic parameters of the Ig4 WT, Ig4 PaTu2 and Ig3-4 WT from urea and thermal denaturation curves calculated using two-state unfolding model.

**Figure 5.**
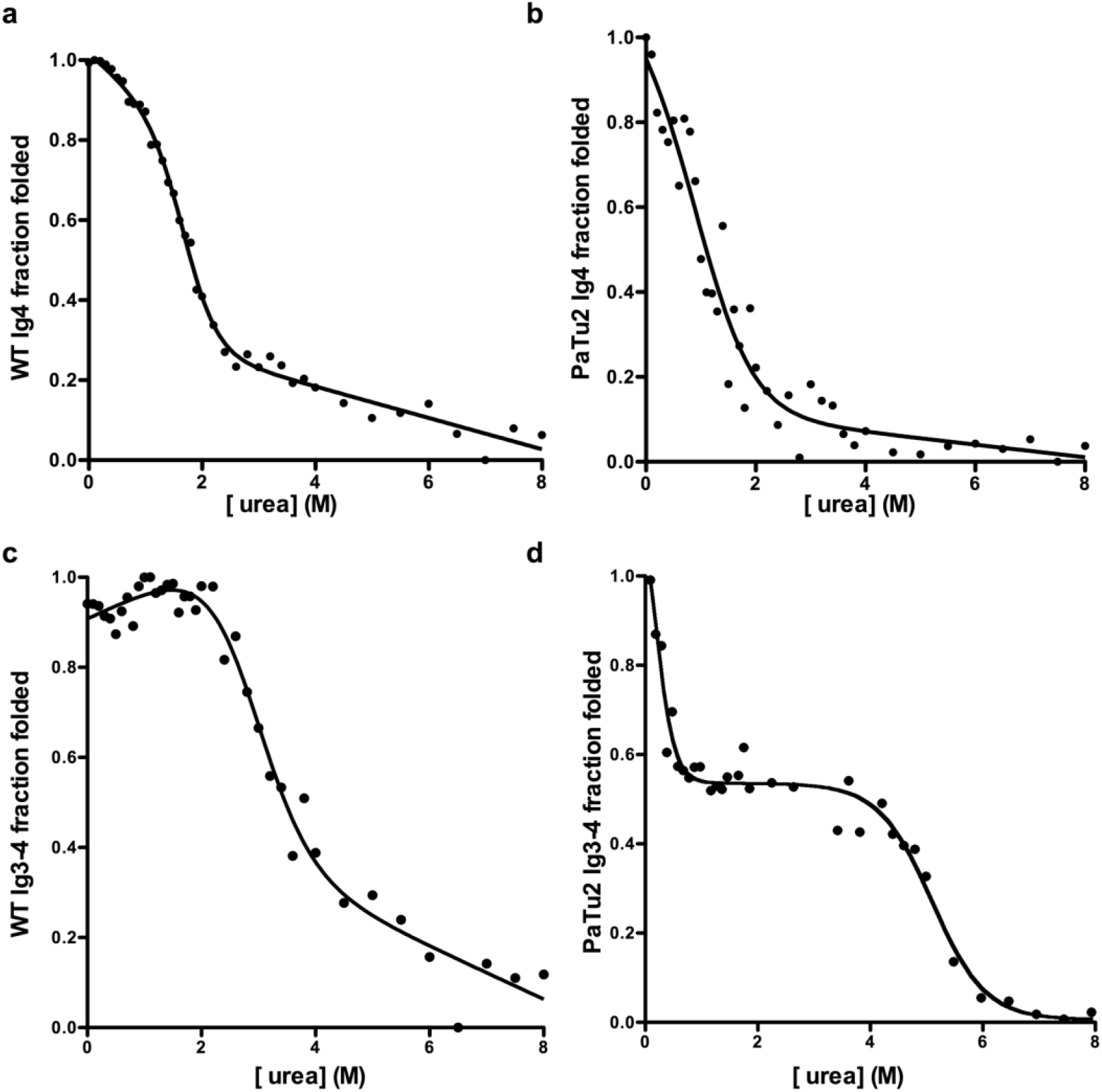
Comparison of palladin domain stability using urea denaturation. Tyrosine emission spectra (Ex: 275 nm, Em: 280-400 nm) were collected for the palladin Ig domains (5 μM) with increasing concentrations of urea. The fraction folded was plotted as a function of urea concentration and data were fitted using equation 2 or 3. Panels correspond to the following proteins: (a) WT Ig4, (b) PaTu2 Ig4, (c) WT Ig3-4, (d) PaTu2 Ig3-4.

Thermal denaturation unfolding curves were also obtained to analyze the change in stability between the WT and PaTu2 Ig4 domains as shown in Figure 6. The melting temperature (T_m_) decreased from 325.5 K in the WT domain to 307.4 K in the PaTu2 mutant domain. The minimal pre-transitional region in the curve also supports the idea that the PaTu2 Ig4 domain is likely partially unfolded even at room temperature. Denaturation of WT Ig4 was nearly reversible while PaTu2 Ig4 was not reversible (Figure S1), reinforcing the importance of the conserved tryptophan for proper folding of the domain. As previously noted for the urea denaturation experiments, two transitions were also observed in the PaTu2 Ig3-4 thermal denaturation spectra as opposed to the singular transition of the WT Ig3-4 spectra.

**Figure 6.**
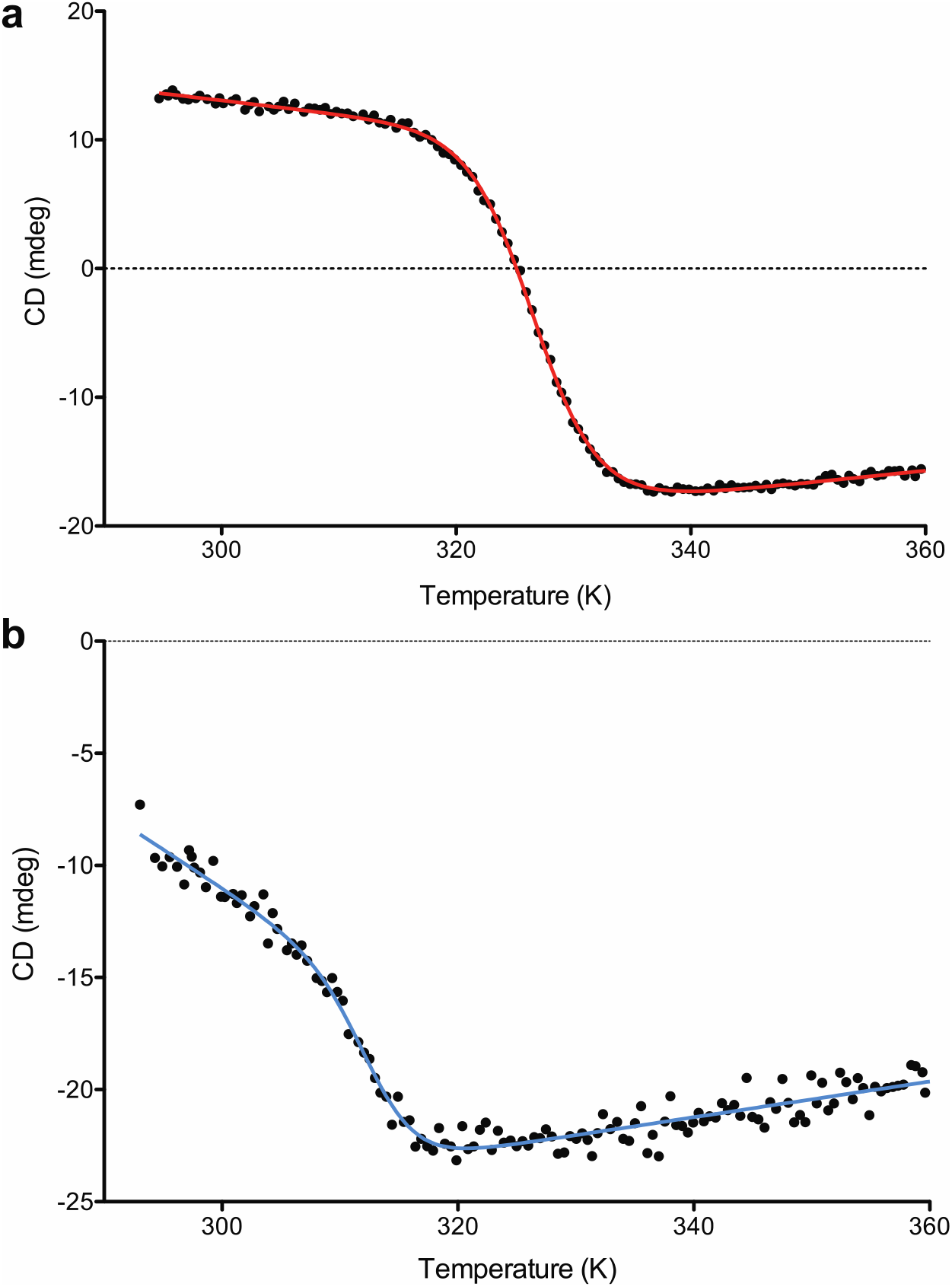
Thermal denaturation curves as monitored by circular dichroism and fitted according to equation 1 for (a) WT Ig4 (b) PaTu2 Ig4.

### 2.5 Fluorescence spectra reveal anomalous tyrosinate emission

When comparing the fluorescence emission spectra of WT Ig4 (A), PaTu2 Ig4 (B), WT Ig34 (C) and PaTu2 Ig34 (D) under native and denaturing conditions in Figure 7, we noticed that all four proteins display similar fluorescence spectra with the exception of PaTu2 Ig4 which does not display the 3 nm red shift between the native and denatured states. A red shift is most likely caused by the unfolding of each Ig domain in 8 M urea, We speculate that the failure to observe a red shift with PaTu2 Ig4 is that this mutant starts off in a partially unfolded state due to the loss of the conserved tryptophan. On the other hand, the red shift observed in the case of PaTu2 Ig3-4 could be attributed to unfolding of the Ig3 domain. These results support the hypothesis that this tryptophan is critical for proper folding of Ig4 domain.

**Figure 7.**
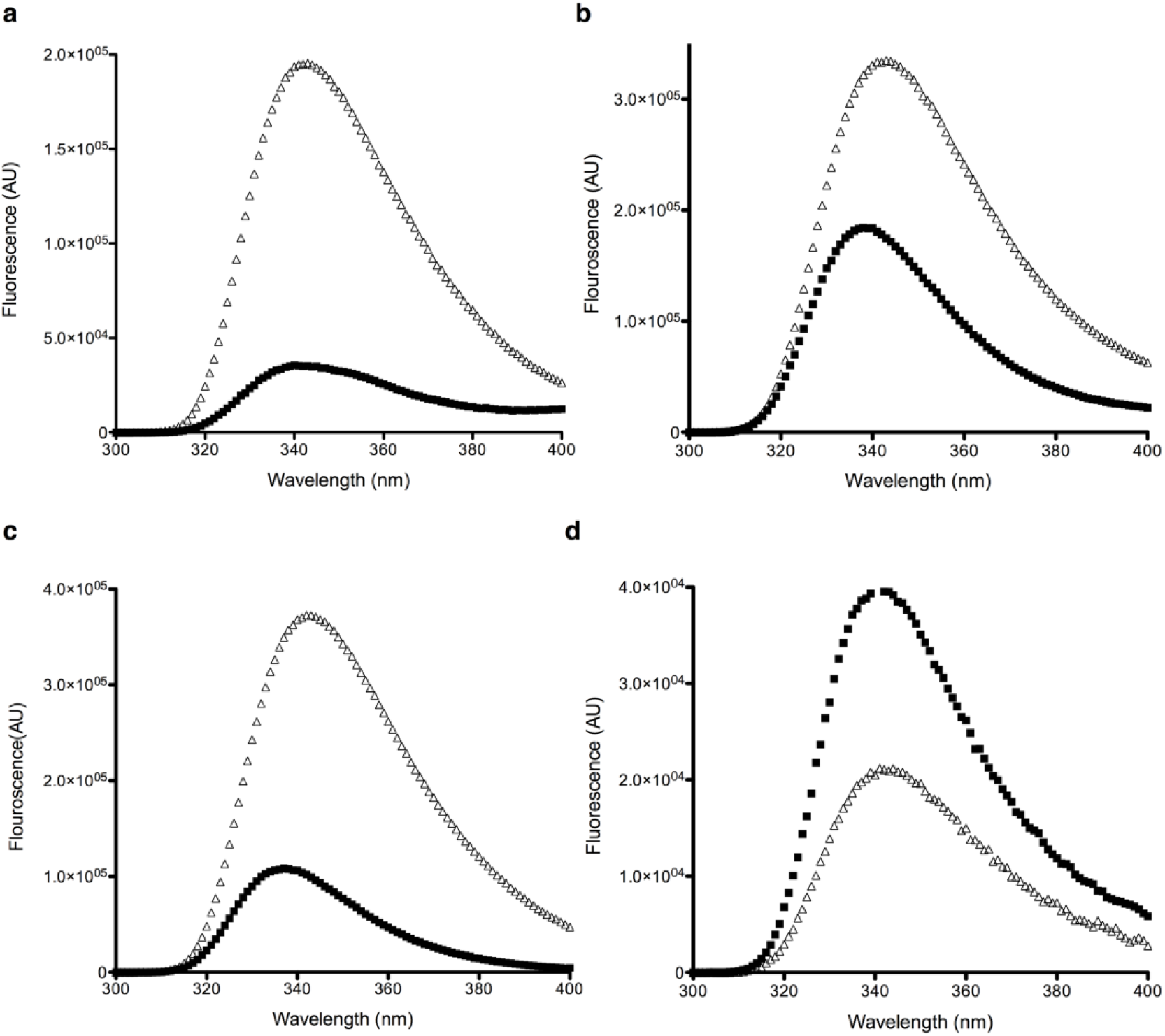
Fluorescence spectra of palladin Ig domains. Emission spectra with 0 M (black squares) or 8 M (white triangles) urea: (a) WT Ig4, (b) PaTu2 Ig4, (c) WT Ig3-4, (d) PaTu2 Ig3-4.

Quantum yield values for these fluorescence spectra were calculated using equation 4 with tryptophan as a reference (Table S1). An increase in quantum yield is seen under chemical denaturing conditions for both WT and PaTu2 Ig4 domains. Conversely, a decrease in quantum yield is observed in PaTu2 Ig4 from pH 6.2 to pH 11 as opposed to the increase in WT Ig4. Most surprising is the observation of a fluorescence maxima at 342 nm for PaTu2 Ig4, a protein domain that contains no tryptophan residues. When we varied the excitation wavelength (Figure S2), the peak height in the emission curve varied slightly; however, the peak maxima remained at 342 nm. This anomalous fluorescence peak has previously been observed in other Class A or tryptophan-free proteins and has been attributed to arise from an excited-state tyrosinate produced either by excited-state proton transfer to adjacent acceptor groups or direct excitation of ground-state tyrosinate.^9–17^

To distinguish between which of these two mechanisms dominates in our case, we studied the influence of pH on the fluorescence of both WT and PaTu2 Ig4 domains. Figure 8 shows the emission surfaces of WT and PaTu2 Ig4, obtained by excitation at 278 nm, over the pH range 6.5 to 11. We observed that the emission at 342 nm was independent of pH below the pKa of the ground-state tyrosine in the PaTu2 Ig4 domain (Figure 8b); therefore, we can surmise that the anomalous tyrosinate peak is most likely due to transfer of an excited-state proton from singlet excited tyrosine based on previous work.^11; 13; 16^ This mechanism is further supported by fluorescence emission spectra of PaTu2 Ig4 at pH 1-2 (Figure S3). Tyrosinate formation is prevented at this low pH as it is well below the pKa of the tyrosine hydroxyl group at 10.5. This ensured protonation of the tyrosine and the fluorescence spectrum peak revealed a 3 nm blue shift from the pH 7.4 spectrum maximum.

**Figure 8.**
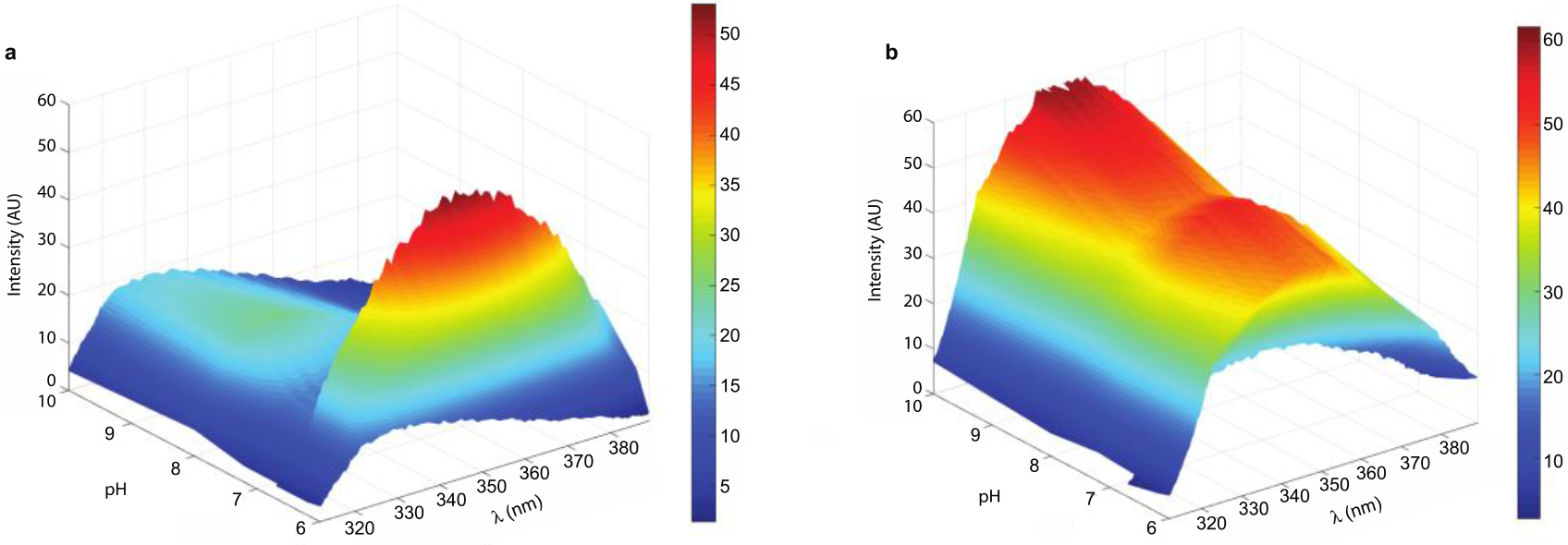
Fluorescence emission scans collected at various pH for (a) WT Ig4 (b) PaTu2 Ig4.

## 3 DISCUSSION

The folding mechanism of immunoglobulin domains has been the subject of numerous studies in several proteins.^18–21^ These studies have established that the hydrophobic core of Ig domains that contain an I-type fold preserve key interactions required for the folding nucleus. While the amino acid composition of the residues that surround the hydrophobic core vary significantly, the invariant positions are occupied by two aromatic amino acids (tryptophan and tyrosine) in the center of the core or “pin region.” Both the position of these amino acids in the pin region and the side chain conformation of these two residues is highly conserved.^1; 22^ Although only the tryptophan is absolutely conserved in all Ig domain containing proteins, the tyrosine is highly conserved among most I-set fold Ig domain proteins such as telokin, titin M5, VCAMD1, and twitchin 39.^22^ Furthermore, the hydrophobic core plays a crucial role in folding, stability, and function of these proteins.

### 3.1 PaTu2 mutation has implications in palladin’s role in crosslinking actin

In this study, we have demonstrated that the conserved tryptophan in the hydrophobic core of palladin’s Ig4 domain plays a critical role in structure, function, and stability. Previous *in vivo* studies identified a single point mutation of the conserved tryptophan in the Ig4 domain of 90-kDa palladin expressed in a pancreatic tumor cell line (PaTu2), and this mutant form of palladin was shown to affect actin bundle formation in these cells.^8^ Previous work also established that the Ig4 domain of palladin is not directly involved in binding or bundling of F-actin; however, the tandem Ig3-4 domain has increased actin-binding affinity as compared to the isolated Ig3 domain.^5^ Furthermore, the palladin Ig3-4 tandem domain also shows increased actin bundling activity as compared to Ig3.^5^ To understand the role of this mutation in actin-binding and bundling by palladin, we made a tryptophan to cysteine substitution mutation in the Ig3-4 domain to mirror the pancreatic tumor mutation. Comparisons of PaTu2 Ig3-4 actin co-sedimentation results with WT revealed a significant difference in actin bundling, but not binding. This is consistent with previous results where the Ig4 domain contributes to actin bundling to a greater degree and this also explains the *in vivo* results where formation of actin bundles is reduced when the PaTu2 mutant form of palladin is expressed in cells. Yet, increasing the concentration of PaTu2 Ig3-4 was able to overcome this deficiency in actin bundle formation and result in crosslinking similar to WT Ig3-4. Additionally, actin binding and bundling is not abolished completely when the PaTu2 mutation is introduced due to the presence of an intact principal actin-binding site that is Ig3 and linker region between two domains present in both the WT and mutant. This suggests that even a misfolded Ig4 domain and/or contributions from the linker region may play a part in increasing actin crosslinking by palladin. Previous work by Dixon *et al.* ruled out any role for the linker as assays with the Ig3 domain plus linker residues (Ig3L) showed no enhancement in binding or bundling above that of the Ig3 domain alone.^5^ Further investigations of the linker residues between Ig3 and Ig4 domains of palladin are warranted, especially in light of the fact that this linker is quite long when compared to other tandem Ig domains. Our structural studies suggest that there is a significant difference in structure between the isolated WT and PaTu2 Ig4 domains and that difference is amplified in the tandem Ig3-4 domain. This suggests that this tryptophan is critical for domain folding and affects both structure and function that may also extend to domain-domain interactions.

### 3.2 Importance of conserved tryptophan in Ig domain structure and stability

Recent NMR backbone dynamics measurements showed that the Ig1 domain residues in the core of myotilin undergo millisecond conformational exchange and the authors suggest that the conserved tryptophan (W283) is likely responsible for maintaining the structural integrity of the entire protein.^23^ Interestingly, this tryptophan in the C strand of the myotilin Ig1 domain is absolutely conserved across all of the Ig domains in the palladin family. One interesting feature observed in the WT Ig4 NMR structure of palladin (PDB: 2DM3) is that both of the hydrophobic core residues tryptophan and tyrosine are in the β-bridge, in contrast to palladin Ig3 and myotilin Ig1 domain where those residues are found in the C-strand of the β-sheet. Studies have shown that the folding transition state of Ig domain containing proteins is stabilized by interactions in the center of the hydrophobic core.^24^ This suggests that palladin family Ig domains also share a similar folding nucleus even though there is a significant difference in amino acid sequence.

Our NMR spectroscopy results indicate that the PaTu2 mutation affects folding and suggests that the conserved core tryptophan is likely involved in the folding nucleus formation, as this mutation fails to induce correct folding of the Ig4 domain. This conclusion is also supported by the observation made during protein overexpression where the Ig4 PaTu2 required an additional tag for solubility that was not required for WT Ig4. This partial unfolding of PaTu2 Ig4 is further confirmed by the results from Ig4 urea denaturation monitored by NMR at 4 M and 8 M urea where neither of the denatured spectra of WT Ig4 overlaps with the PaTu2 Ig4 spectra.

In a recent study, J. Clark *et al.* observed significant differences in the stability of proteins with Ig domains and provided evidence that this difference is largely due to variations in the rate of unfolding.^24^ In our urea denaturation results shown in Figure 5, the PaTu2 mutation resulted in decreased stability as compared to WT. Previous studies indicated that strands B–C and E–F form the folding core during initial stages of Ig domain folding, while both the N- and C-termini fold in later stages and conversely during unfolding, the termini regions unfold early.^18; 20; 25^ Insertion of amino acids to extend the loop length near the folding nucleus decreases stability in the titin 27^th^ Ig domain.^26^ From these observations, one can conclude that a decrease in the stability of Ig domains containing a tryptophan mutation is likely due to failure to form the core folding nucleus structure and subsequent stability contribution results from a partially folded structure.

### 3.3 Anomalous tyrosine fluorescence indicates formation of tyrosinate

While fluorescence intensity above 310 nm is usually attributed to tryptophan residues^12^, this is clearly not the case with the peak observed at 342 nm in our tryptophan-free PaTu2 mutant Ig domain (Fig. 7B). Our fluorescence spectra (Figures 7 and 8) and quantum yield analysis (Table 1) also indicate that this fluorescence is likely from transfer of an excited state proton from singlet excited tyrosine. The anomalous tyrosinate fluorescence phenomenon has only been reported once in four decades.^27^ While there has been a lot of speculation regarding the conditions that promote the formation of tyrosinate, conclusive studies have yet to be carried out to distinguish between excited state protolysis or direct excitation of ground state tyrosinate. Future studies that systematically probe the residues and structural environment in a variety of proteins displaying tyrosinate fluorescence will hopefully allow us to elucidate the mechanism of tyrosinate formation. This insight will help to identify proteins with similar spatial amino acid arrangements and allow us to determine the prevalence of tyrosinate fluorescence phenomena that may be being overlooked in the interpretation of protein fluorescence spectra. For instance, fluorescence intensity in the 330-350 nm range is often used to monitor conformational and structural perturbations in proteins; however, this intensity could be misinterpreted in proteins that form tyrosinate residues at unexpected pH values.

## 4 CONCLUSION

These results suggest that the Ig3-4 domain structure is critical for actin bundling and changes in Ig4 structure result in decreased bundling even though this domain is not directly involved in binding actin. This observation is consistent with our recent study where we showed the inter-domain flexibility and conformation changes in the actin bound and free form of palladin Ig3-4 are critical for actin bundling.^6^ From these results, we hypothesize that orientation of the Ig4 domain with respect to the Ig3 domain in the PaTu2 Ig3-4 domain is altered by the PaTu2 mutation, which has significantly disturbed the Ig4 domain structure, thereby decreasing actin bundling. This study also emphasizes the critical role of the tryptophan that is absolutely conserved across all palladin family member Ig domains and plays an important role in domain structure, folding, stability, and function.

## 5 MATERIALS AND METHODS

### 5.1 Expression and purification of palladin Ig domains

The *Mus musculus* Ig3, Ig4, and Ig3-4 domains of palladin were purified as described previously.^28^ Initial attempts to express and purify PaTu2 Ig4 from the pTBSG vector were unsuccessful as the protein are sequestered in inclusion bodies. To overcome this problem, we subcloned both the wildtype and PaTu2 Ig3-4 sequences into pTBMalE expression vector, which includes both a His_6_ and MBP solubility tag to maintain Ig3-4 in the soluble fraction. Likewise, we have subcloned the PaTu2 Ig4 into the pTBMalE expression vector as described.^6^ All constructs were transformed into BL21 (λ DE3) *Escherichia coli* cells for protein expression and cell cultures were grown at 37 °C until the OD_600_ reached 0.7 and were further grown overnight at 18 °C in ZYM-5052 auto-induction media as described.^29^ Cell cultures were pelleted down and resuspended into lysis buffer (20 mM PIPES, pH 8.0; 300 mM NaCl and 10 mM imidazole), followed by sonication and clearing of cell lysate by centrifugation. All the constructs were purified on a Ni-NTA column according to the manufacturer’s guidelines (ThermoFisher Scientific; Waltham, MA). The N-terminal histidine tag and MBP tag were cleaved by TEV protease and further purified as previously reported.^5^ All palladin domains were finally dialyzed into HBS buffer (20 mM Hepes, pH 7.5; 100 mM NaCl and 1 mM DTT).

Protein concentration was determined using the bicinchoninic acid (BCA) assay as determination of protein concentration by measurement of absorbance at 280 nm relies on the presence of tryptophan (Pierce™ BCA Protein Assay Kit-Reducing Agent Compatible; ThermoFisher Scientific; Waltham, MA).

### 5.2 Actin co-sedimentation assays

Actin was purified from rabbit muscle acetone powder as described previously.^30^ Purified G-actin was diluted into 2X polymerization buffer (10 mM Hepes, pH 7.5; 100 mM KCl and 2 mM MgCl_2_) for 30 min to make F-actin. In the actin-binding assay, palladin Ig domains (5 μM) were incubated with varying concentration of F-actin (0–20 μM) for 1 hr followed by high-speed centrifugation at 100,000 xg for 30 min to separate supernatant and pellet. In the actin bundling assay, F-actin (10 μM) was incubated with varying concentrations (0–40 μM) of palladin Ig domains for 1 hr, followed by low speed centrifugation at 5,000 xg for 10 min to separate bundles (B). A subsequent high-speed centrifugation step (100,000 xg for 30 min) was then carried out to separate actin in the supernatant (S) and pellet (P). In both assays, the bundle, pellet and supernatant fractions were resuspended in 2X SDS buffer and samples were separated by SDS-PAGE. In the bundling assay, actin, and in the binding assay, palladin Ig-domains, are quantified by using ImageJ software.^31^

### 5.3 Circular dichroism

CD spectra were recorded using a Jasco J810 spectropolarimeter. Each Ig domain was extensively dialyzed into 10 mM phosphate buffer, 50 mM sodium fluoride, 1 mM TCEP, pH 7.5. Spectra were recorded on 20 μM of protein between 190 nm and 260 nm in a 0.1 mm cell at room temperature with the 10 nm/min scan speed. Thermal denaturation unfolding curves were recorded for both WT and PaTu2 mutant of Ig4. Spectra were recorded on 20 μM of protein at 206 nm in a 0.1 mm water-jacketed cell that was heated from 293.15 K to 338.15K as controlled by a JULABO water circulator. This wavelength was chosen because it showed the greatest difference between the native and unfolded samples. To examine reversibility, the sample was then cooled from 338.15 K to 293.15 K In addition, one sample was melted at 338.15 K and a wavelength spectrum was obtained from 198 nm to 260 nm. The sample was then left at 277.15 K overnight and another spectrum was collected. Thermal denaturation under these conditions was 82% reversible for WT Ig4. The enthalpy of unfolding (ΔH_m_°) and the midpoint thermal transition (T_m_) were determined by a least-squares fit of the data to a modified van’t Hoff equation describing a two-state unfolding reaction using the following equation with GraphPad Prism software:

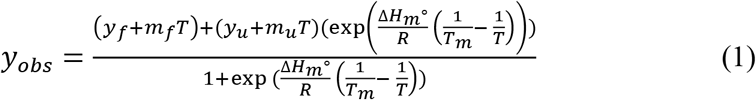

where the parameters y_f_, m_f_, y_u_, and m_u_ refer to the y-intercept and slope of the pre- and post-unfolding baselines, respectively. R is the gas constant (1.987 cal/mol*deg)) and T is the absolute temperature (in Kelvin).^32^

### 5.4 Fluorescence spectroscopy

Fluorescence spectra of both WT and PaTu2 mutants of Palld-Ig3, Ig4, and Ig3-4 domains were collected on a PTI spectrofluorometer with excitation at 275 nm and emission at 280-400 nm. For urea denaturation experiments, samples were prepared with 5 μM palladin Ig domains in 20 mM Hepes pH 7.5, 100 mM NaCl with gradual increase in the concentration of urea from 0 to 8 M. Samples were incubated overnight and tyrosine emission spectra collected. The fraction folded was calculated from emission spectra peak maxima and plotted against urea concentration. The data was then fit to the two state (N ↔ U) equation 2 or three state (N ↔ I ↔ U) equation 3 with GraphPad Prism software:

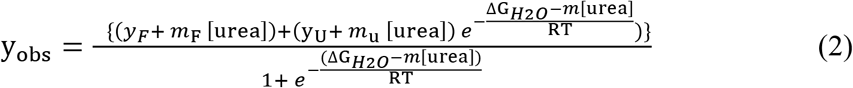

where y_F_ and y_U_ are intercepts and mF and mU are slopes of folded and unfolded state respectively and m is the measure of dependence of ΔG_0_ on the denaturant concentration.^33; 34^

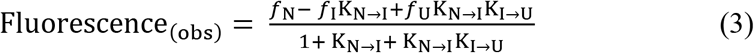

where *f*_N_, *f*_I_, and *f*_U_ are fraction of native, intermediate and unfolded state and K_N→I_ and K_I→U_ are equilibrium constants obtained as described previously.^34^

Quantum yield was calculated from fluorescence spectra with 20 μM protein samples with excitation at 275 nm and emission from 300-700 nm according to:

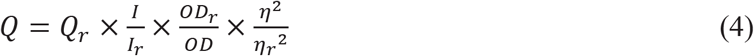

where Q and Q_r_ are the quantum yields for the sample and the reference, respectively. I and I_r_ stand for the total fluorescence as determined from the area under the emission spectra. OD and OD_r_ are the optical density at the excitation wavelength while η and η_r_ are the refractive index of the sample solvent.

## ABBREVIATIONS

Ig: immunoglobulin
NMR: nuclear magnetic resonance
PaTu2: pancreatic tumor cell line and also used to refer to specific W40C mutation found in palladin in this cell line
WT: wildtype
CD: circular dichroism
MBP: maltose binding protein
HSQC: heteronuclear single quantum correlation spectroscopy
C_m_: folding transition mid-points
PDB: protein database
OD_600_: optical density at 600 nm
His_6_: six histidine protein tag
Ni-NTA: nickel nitrilotriacetic acid
DTT: dithiothreitol
SDS-PAGE: sodium dodecyl sulfate polyacrylamide gel electrophoresis

## SUPPORTING MATERIAL

Supplementary material contains Figures S1, S2, S3 and Table S1.

## ACKNOWLEDGEMENTS

SVS and JTD were supported by K-INBRE (P20GM103418) and WSU undergraduate research grants. SVS was also supported by the McNair Scholars program at WSU. RAK was supported by the Erach R. Talaty Graduate Research Assistantship. AA was supported by a Watkins Summer Research Participation Fellowship. This research was supported by an Institutional Development Award (IDeA) from the National Institute of General Medical Sciences of the National Institutes of Health under grant number P20GM103418 and P30 GM1110761 as well as an R15 with Award Number R15GM120670 awarded to MRB. The authors thank members of the Beck lab for many helpful discussions, and Carol Otey and Dan Arneman for initial experiments with the PaTu2 mutant in cells.

