## Supporting Material for "Conserved tryptophan mutation disrupts structure and function of immunoglobulin domain revealing unusual tyrosine fluorescence"

### SUPPLEMENTAL INFORMATION

**Figure S1.** Thermal denaturation and reverse spectra of (a) WT Ig4 and (b) PaTu2 Ig4. Voltage readings for data in (a) and (b) shows no sign of aggregation.

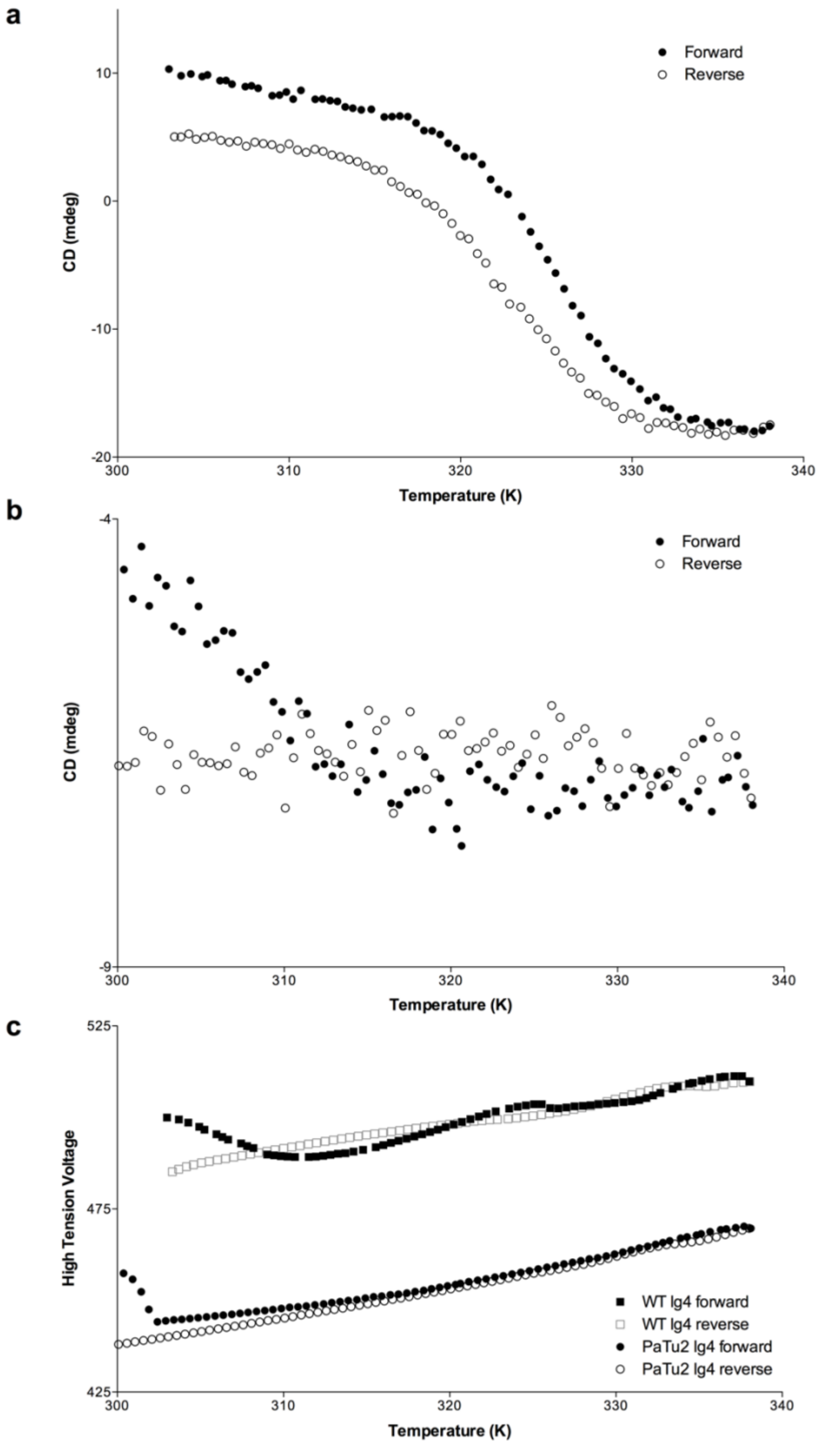

**Figure S2**. Comparison of PaTu2 Ig4 fluorescence emission spectra produced from varying excitation wavelengths (260, 275, 280, or 290 nm).

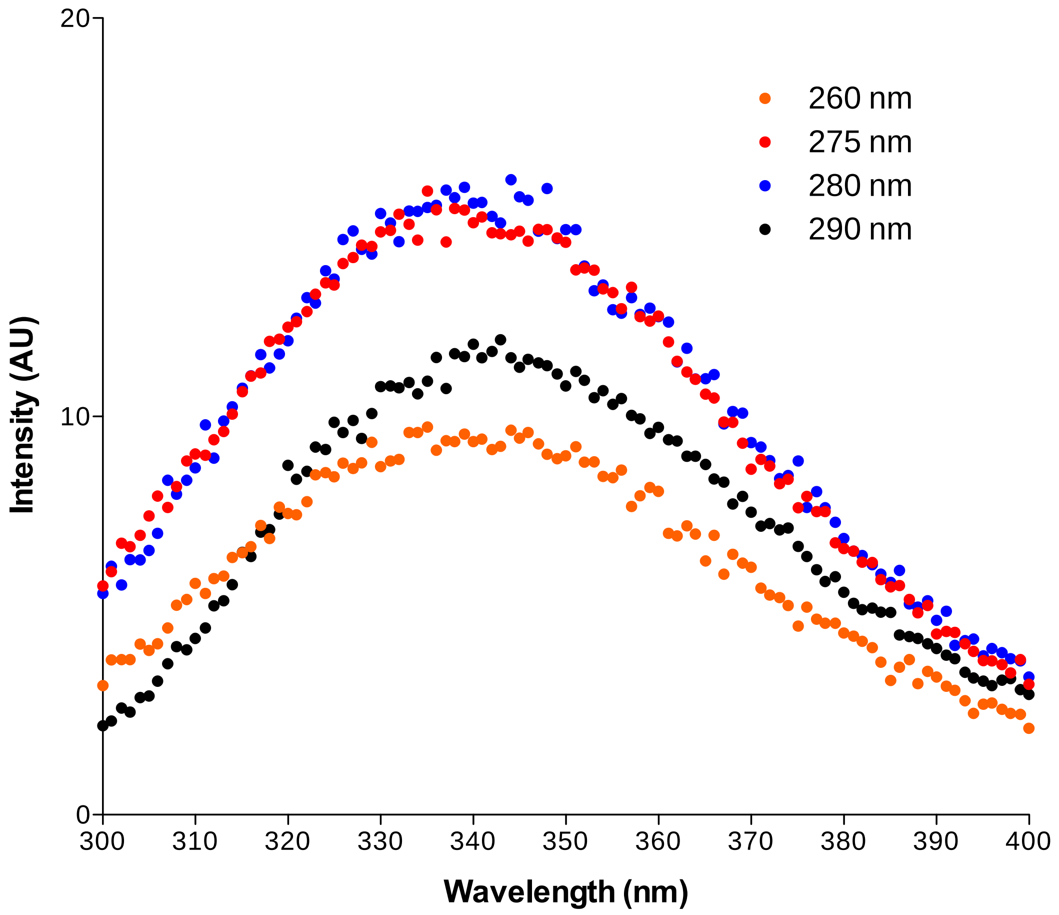

**Figure S3.** Fluorescence emission spectra of Ig4 domains at physiological pH and very acidic pH.

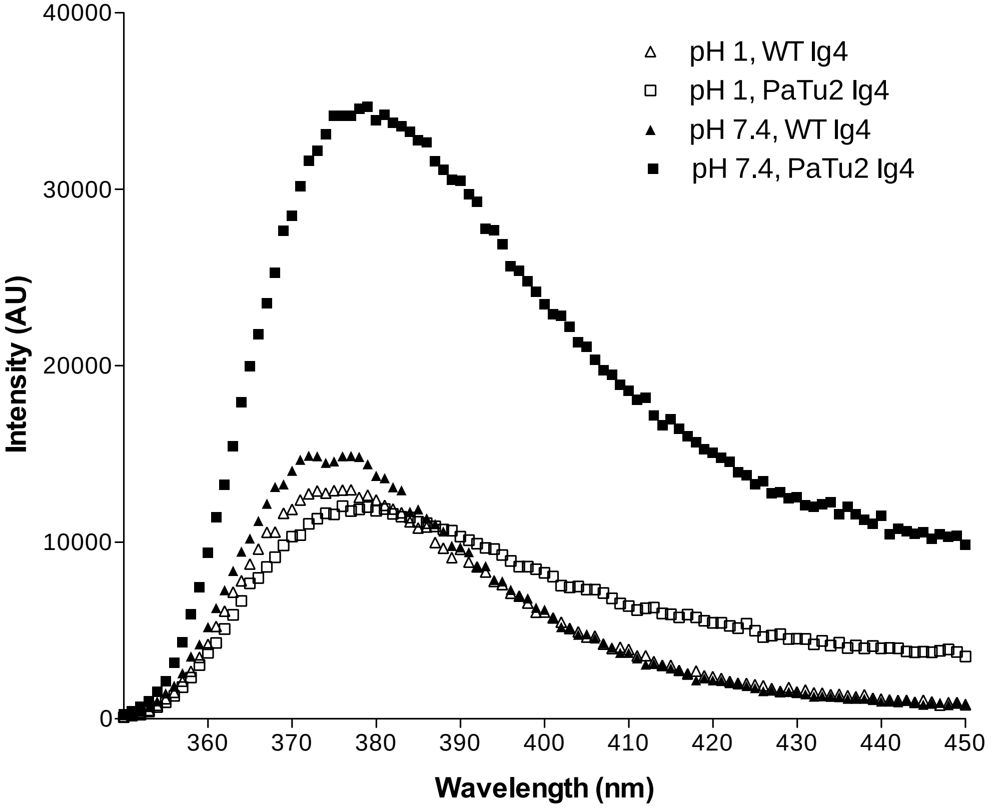

**Table S1. Quantum yields from fluorescence spectra using tryptophan as reference**

| | WT Ig4 | | | | PaTu2 Ig4 | | | |
|---|---|---|---|---|---|---|---|---|
| | 0 M urea | 8 M urea | pH 6.2 | pH 11 | 0 M urea | 8 M urea | pH 6.2 | pH11 |
| Quantum Yield | 0.008 | 0.104 | 0.013 | 0.071 | 0.013 | 0.044 | 0.039 | 0.026 |
